# RNALens: Study on 5’ UTR Modeling and Cell-Specificity

**DOI:** 10.1101/2025.07.20.665722

**Authors:** Lei Mao, Yuanhe Tian, Kang-wei Qian, Yan Song

## Abstract

Recently, the Transformer architecture has been applied to predict the structure, function, and regulatory activity of biological sequences. Predicting the cell-specific regulatory impact of 5’ untranslated regions (5’ UTRs) on mRNA expression and translation remains a key challenge for rational mRNA design. Existing studies such as UTR-LM, RNABERT, and RNA-FM train transformer-based models solely on 5’ UTR sequences with fixed nucleotide tokenization schemes and auxiliary structural features. These models pay less attention to the integration of broader genomic context and thermodynamic objectives, which limits their ability to generalize across diverse cell types and accurately predict both mRNA expression level (EL) and translation efficiency (TE). In this paper, we propose RNALens, a foundation model pre-trained in two stages on multispecies genomic sequences and curated 5’ UTR data using masked language modeling augmented with secondary structure prediction and minimum free energy regression. RNALens employs byte-pair encoding to capture variable-length nucleotide motifs. It is then fine-tuned on high-throughput reporter assay datasets from HEK293T, PC3, and muscle tissues to yield specialized predictors for EL and TE in each cellular context. Experiment results on benchmark datasets demonstrate that RNALens achieves superior performance than existing machine learning methods for both expression and translation predictions across cell-specific and cross-context tests, offering an efficient in silico platform for guiding the design of mRNA therapeutics with precise cellular targeting.^1^

## 1 Introduction

As mRNA therapeutics expand to more complex disease applications such as targeted protein replacement therapies for specific organs [1–3], therapeutic vaccines targeting specific cancer cells [4–6], or gene editing [7, 8], the need for high-precision targeted delivery and cell- or tissue-specific expression become increasingly critical. Achieving cell- or tissue-specific protein expression is a core strategy for enhancing the safety and efficacy of mRNA drugs [9–11]. Such precise regulation helps prevent adverse reactions in non-target cells, improving the overall safety and tolerability of the drug [9–11]. In an mRNA molecule, the 5’ untranslated region (5’ UTR) is a non-coding sequence located between the transcription start site and the translation start codon, and plays a crucial role in gene expression regulation [12, 13]. The 5’ UTR is considered a “post-transcriptional regulatory hub” that influences mRNA stability, splicing, translation efficiency, and subcellular localization [12, 14–16]. 5’ UTR serves as a binding site for various RNA-binding proteins (RBPs) and trans-acting factors [15], such as non-coding RNAs (ncRNAs) [17–20]. The expression of these RBPs [21–23] and ncRNAs [24–26] often exhibits cell-type specificity, indicating that their interactions with the 5’ UTR serve as a crucial endogenous mechanism for achieving precise cell- or tissue-specific mRNA regulation in organisms. In other words, a certain cell type harbors a distinct intracellular environment that influences mRNA stability and translation through modulation of 5’ UTR regions. Therefore, this region integrates various intracellular and extracellular signals to fine-tune gene expression according to specific cellular needs and environmental changes, making it an ideal target for engineering desired expression profiles. The development of high-throughput screening technologies, such as massively parallel reporter assay (MPRA) [27], significantly advances the screening of mRNA sequences, including 5’ UTR [28]. However, to identify expression specificity across different cell types, one necessarily requires the same set of 5’ UTR sequences to exist across these cell types, which is challenging to execute at the throughput scale comparable to MPRA’s.

In recent years, large language models (LLMs), particularly those based on the Transformer architecture, achieve revolutionary breakthroughs in natural language processing [29–32] and are rapidly applied to the analysis of biological sequences (e.g., DNA, RNA, and protein) to predict their structure, function, and regulatory activity [33–39]. LLMs are able to learn complex patterns, contextual information, and long-range dependencies from vast amounts of sequence data, demonstrating potential superior to traditional machine learning methods in genomic and transcriptomic analysis [35, 36]. They interpret the “language” of genetic information with unprecedented scale and accuracy [35, 36]. Such foundation models are fine-tuned to execute multiple downstream tasks with relatively small-scale training data and achieve better performance compared to models trained from scratch [35–37, 39–41]. Currently, a series of RNA LLMs have emerged, such as UTR-LM [41], RNABERT [33], RNA-FM [42], BigRNA [43], and LAMAR [44], which show application prospects in RNA structure prediction, functional annotation, and regulatory activity prediction. Therefore, it is expected to apply LLM techniques to modeling 5’ UTR. Among these RNA LLMs, UTR-LM [41] is one of the most representative models to predict the effect of 5’ UTR on the protein expression level. UTR-LM uses four single nucleotides as tokens, which limits its ability to represent UTR sequences due to a low vocabulary size, and it is only pre-trained on UTR sequences with-out more general genome contexts, which prevents it from being generalized to more cases. Therefore, it is expected to build a powerful foundation model for 5’ UTR with advanced tokenization approaches and trained on more data.

In this paper, we propose a novel RNA foundation model named RNALens, which employs a standard Transformer encoder architecture. RNALens is pre-trained on both genomic sequences data and 5’ UTR sequences data, aiming to learn RNA-specific sequence features, such as short-sequence features, regulatory patterns, and long contextual information. RNALens is then fine-tuned to predict the cell- or tissue-specific function of 5’ UTR on mRNA expression level (EL) and translation efficiency (TE). The experiments on benchmark datasets demonstrate the effectiveness of RNALens, which presents superior performance to existing models. In addition, RNALens presents noticeable application values. Since RNALens is fine-tuned on data with different cell or tissue types, it may have the potential to serve as a virtual experiment environment that predicts the response to a certain perturbation in cells. For example, the RNALens model fine-tuned on the sequencing data obtained in HEK293T cell (human embryonic kidney cell line) represents the expression environment in this cell type and is able to predict the expression level of a given 5’ UTR sequence with high accuracy. Therefore, the RNALens model has great application prospects.

## 2 Results

The expression of mRNA is the foundation of protein translation, and both the expression level (EL) and translation efficiency (TE) of mRNA collectively determine the final protein synthesis level. We present the experiment results of RNALens in predicting mRNA EL and TE, and its ability to capture cell-type-specific regulatory signatures for both mRNA EL and TE. We evaluate the performance using the Spearman correlation coefficient between predicted and gold standard values.

### 2.1 RNALens in mRNA Expression Level prediction

The RNALens model utilizes a Transformer encoder architecture. During the pre-training phase, to enable the learning of both general features of RNA se-quences and specific regulatory signals of 5’ UTRs, we use different proportions of genomic sequence data and 5’ UTR sequence data, which results in two foundation models, named RNALens-1 and RNALens-2. This pre-training strategy, combining broad genomic context with specific regulatory region data, aims to endow the model with stronger generalization abilities and a deeper understanding of 5’ UTR functions.

Subsequently, we fine-tuned the two RNALens foundation models, namely, RNALens-1 and RNALens-2, for the prediction of mRNA expression level (EL) by 5’ UTR in different cell or tissue types, using the 5’ UTR library data from Cao et al. [45]^2^ In total, six fine-tuned models are obtained for HEK293T cell, PC3 cell, and muscle tissue, two for each cell or tissue type, based on RNALens-1 and RNALens-2, respectively. In the experiments, EL is measured by RNA-seq RPKM (Reads Per Kilobase per Million mapped reads) [41]. The results of these models in predicting mRNA EL based on the given 5’ UTR sequence are reported in Table 1, and the relation between the predicted value and the gold standard value is reported in Figure 1A and Figure 1B, where Figure 1A reports the results for RNALens-1 and Figure 1B shows the results for RNALens-2.

**Table 1:**
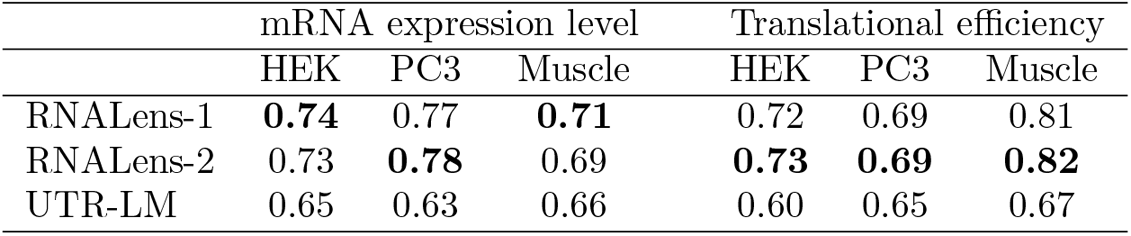
Prediction of EL and TE by RNALens-based models or benchmarks in different cell/tissues types. Spearman’s correlation coefficients between predicted values and true values are presented in this table. The best results are highlighted in boldface.

**Figure 1:**
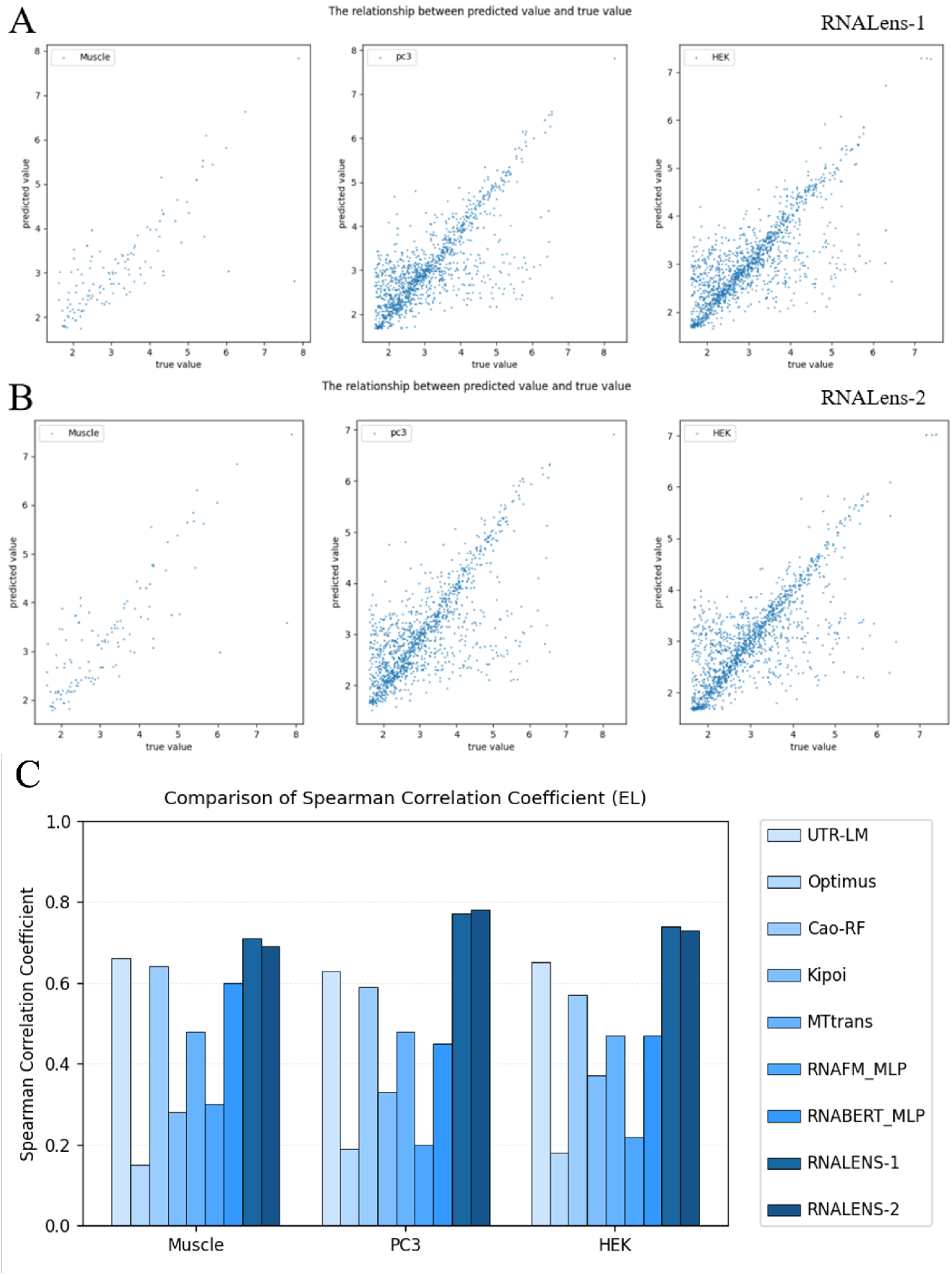
Prediction of mRNA expression level (EL). Scatter plots show the correlation between true mRNA EL value and mRNA EL value in HEK293T cells, PC3 cells or muscle tissues predicted by models fine-tuned from RNALens-1 (A) and RNALens-2 (B). (C) RNALens-based models outperformed on mRNA EL prediction when compared to other benchmarks.

We then compare the performance of our model with a series existing approaches, including Optimus [28], Cao-RF [45], relevant models from the Kipoi model repository [46], MTtrans [41,47], RNAFM MLP [41,42], RNABERT MLP [33, 41], and UTR-LM [41]. Among these models, UTR-LM is a state-of-the-art model that presents strong performance on mRNA EL prediction. Evaluations based on Spearman’s Correlation Coefficient (see Figure 1C and Table 1) indicate that cell- or tissue-specific models fine-tuned from RNALens-1 and RNALens-2 consistently outperform all existing models, including UTR-LM, in predicting 5’ UTR-regulated mRNA ELs in their respective cell or tissue environments. The superior performance of RNALens is attributed to its unique pre-training strategy, which combines broad genomic contextual information with 5’ UTR-specific sequence data. This enables it to learn a deeper, more universal sequence “grammar” and regulatory logic, which is more effectively utilized when fine-tuning for specific cell types. In contrast, although UTR-LM is also designed for 5’ UTRs and incorporates information such as secondary structure and minimum free energy during pre-training [41], it lacks the genome-level pre-training that provides a broader feature learning space.

### 2.2 Models for Predicting Translation Efficiency in Different Cell or Tissue Types

Protein production is not always correlated with mRNA levels [48], as there are numerous translational and post-translational regulatory mechanisms, where some of which are cell- and tissue-specific. As such, translation efficiency (TE) represents an additional layer of regulation that provides important insights beyond mRNA expression levels alone. Notably, 5’ UTRs contain multiple binding sites for RBPs and ncRNAs, which regulate cell- and tissue-specific TE through ribosome recruitment. Since the ribosomal footprint on the mRNA is able to be measured by Ribo-seq, TE is calculated as the ratio of Ribo-seq RPKM to RNA-seq RPKM for a given mRNA. To explore RNALens’s capability in predicting TE, we utilize publicly available Ribo-seq data and RNA-seq data from Cao et al. [45] to perform fine-tuning for RNALens-1 and RNALens-2 in HEK293T cells, PC3 cells, and muscle tissues, respectively, thereby constructing a suite of RNALens-TE models for different cell or tissue types. As shown in Figure 2 and Table 1, the RNALens-TE models exhibit high TE prediction accuracy. Furthermore, RNALens-TE models outperformed all benchmark models tested, including UTR-LM (see Figure 2 and Table 1).

**Figure 2:**
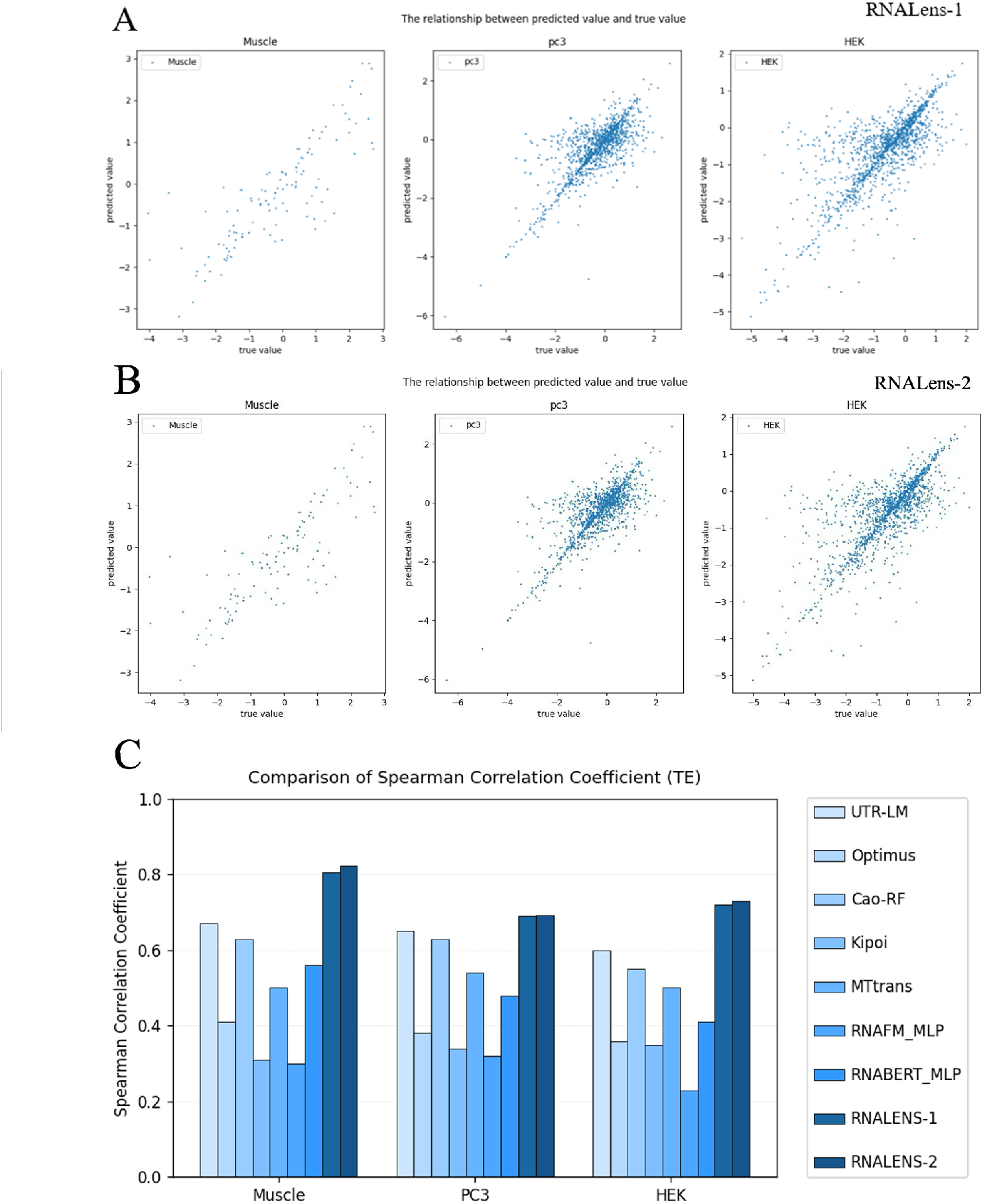
Prediction of mRNA translation efficiency (TE). (A) Scater plots showed the correlation between true mRNA EL value and mRNA EL value in HEK293T cells, PC3 cells or muscle tissues predicted by models fine-tuned from RNALens-1. (B) Scater plots showed the correlation between true mRNA EL value and mRNA EL value in HEK293T cells, PC3 cells or muscle tissues predicted by models fine-tuned from RNALens-2. (C) RNALens-based models outperformed on mRNA TE prediction when compared to other benchmarks.

### 2.3 RNALens Captures Cell-Type Specific Regulatory Signatures

To further investigate whether the RNALens-based models learn cell- and tissue-specific information in mRNA expression regulation, we calculate the Spearman’s correlation coefficient of the mRNA EL value predicted by three models fine-tuned based on RNALens-1, named RNALens-HEK, RNALens-PC3, and RNALens-Muscle, with the EL value measured by RNA-seq in all three cell or tissue types. The results are shown in Figure 3A and Table 2. Consistent with previous findings, each model achieved the highest correlation in its corresponding cell or tissue type (0.7378 for RNALens-HEK, 0.7138 for RNALens-PC3, 0.7728 for RNALens-Muscle). In contrast, cross-context predictions reveal that the correlation between RNALens-HEK predictions in PC3 cells (0.3868) and RNALens-PC3 predictions in HEK293T cells (0.4684) is substantially higher than that observed when either model was applied to muscle tissue (0.0046 for RNALens-HEK and 0.0218 for RNALens-PC3) . Consistently, the expression profiles predicted by the RNALens-muscle model in HEK293T cells (0.0140) and in PC3 cells (0.0680) are relatively low. In addition, models fine-tuned based on RNALens-2 are also validated on cell- and tissue-specific information learning and the results are consistent with those of RNALens-1 (Table 2).

**Table 2:**
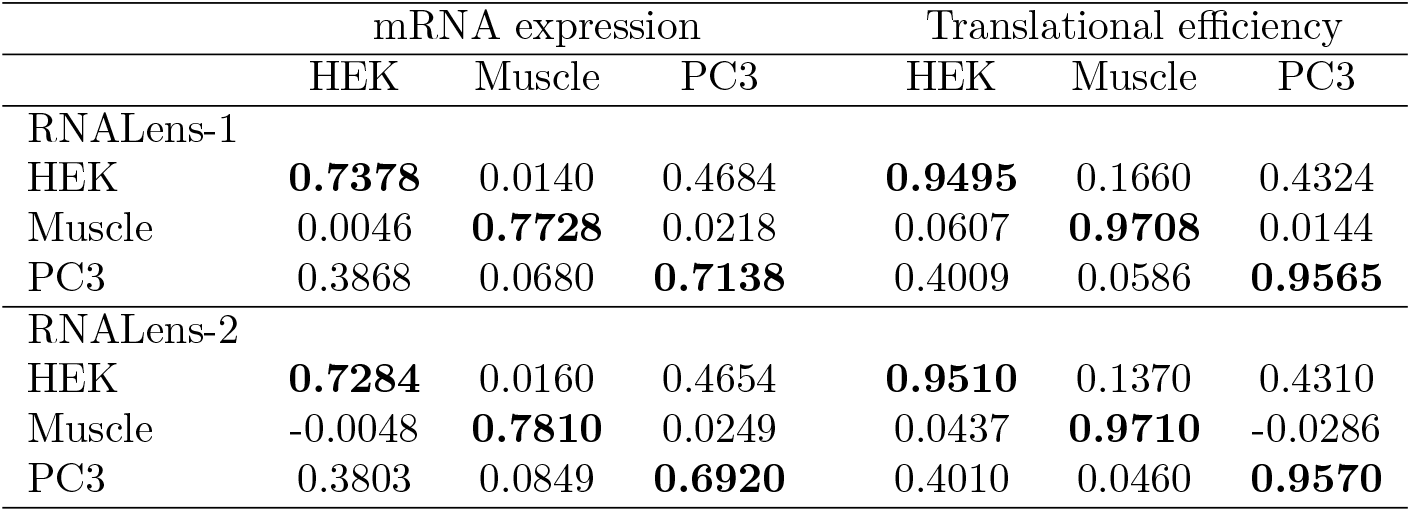
Cell or tissue-specificity in mRNA EL and TE prediction by RNALens-based models. Performance of RNALens-based models fine-tuned on data from HEK293T cells, PC3 cells and muscle tissues are tested in corresponding and non-corresponding cell or tissue contexts. The best results are highlighted in boldface.

|  | mRNA expression |  |  | Translational efficiency |  |  |
| --- | --- | --- | --- | --- | --- | --- |
|  | HEK | Muscle | PC3 | HEK | Muscle | PC3 |
| RNALens-1 |  |  |  |  |  |  |
| HEK | **0.7378** | 0.0140 | 0.4684 | **0.9495** | 0.1660 | 0.4324 |
| Muscle | 0.0046 | **0.7728** | 0.0218 | 0.0607 | **0.9708** | 0.0144 |
| PC3 | 0.3868 | 0.0680 | **0.7138** | 0.4009 | 0.0586 | **0.9565** |
| RNALens-2 |  |  |  |  |  |  |
| HEK | **0.7284** | 0.0160 | 0.4654 | **0.9510** | 0.1370 | 0.4310 |
| Muscle | -0.0048 | **0.7810** | 0.0249 | 0.0437 | **0.9710** | -0.0286 |
| PC3 | 0.3803 | 0.0849 | **0.6920** | 0.4010 | 0.0460 | **0.9570** |

**Figure 3:**
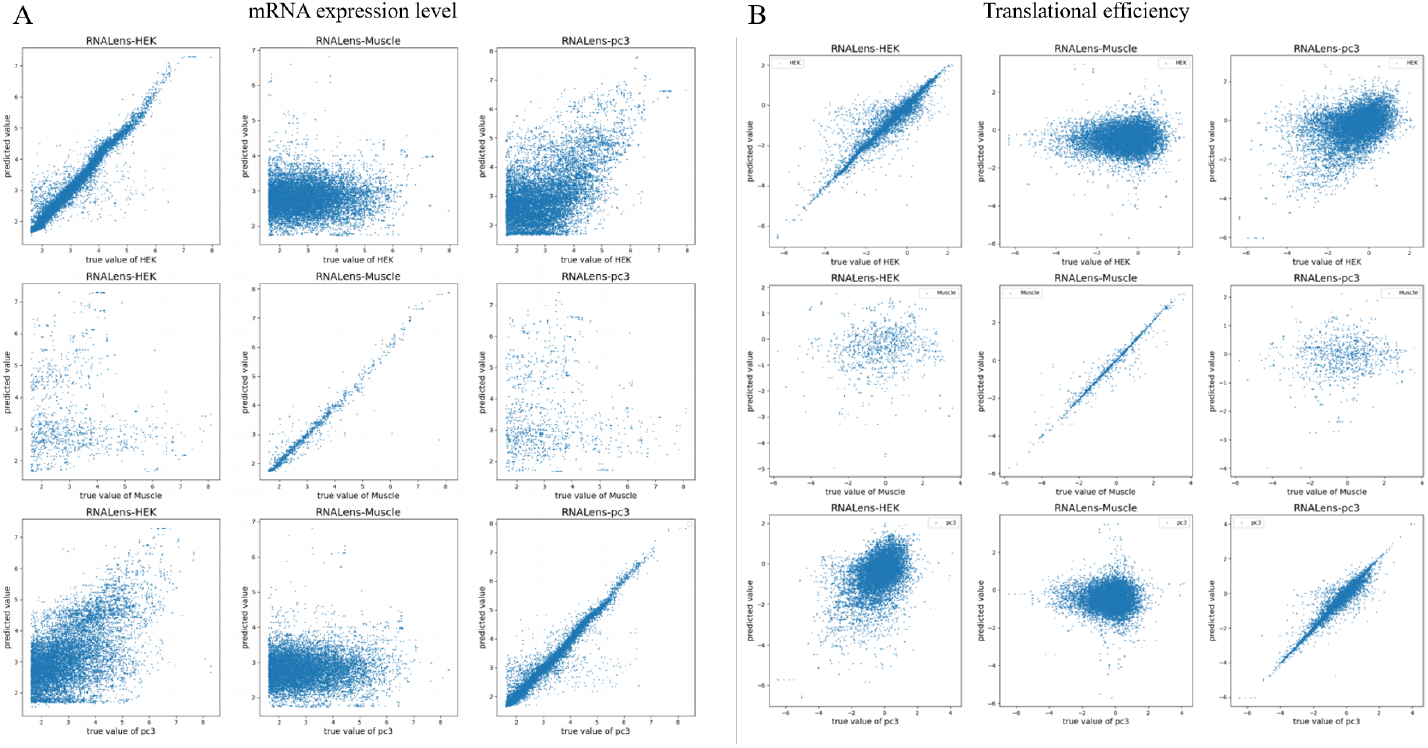
Cell- or tissue-specific prediction of mRNA EL (A) and TE (B) by models fine-tuned from RNALens-1.

We also test RNALens-TE models’ abilities on cell- and tissue-specific information learning, where similar results are observed. The scatter plots of RNALens-1-based models are shown in Figure 3B, and the Spearman’s correlation coefficients of all RNALens-1 and RNALens-2-based models are shown in Table 2.

These results suggest that RNALens learn the cell- or tissue-specific context associated with 5’ UTR function, as well as shared features across related cell types, such as HEK293T cells and PC3 cells, both of which are derived from the urogenital system.

## 3 Discussion

### 3.1 RNALens is a Superior Predictive Tool for Cell- and Tissue-Specific 5’ UTR Function

In this paper, we develop a foundation model named RNALens for RNA sequences. Compared to conventional deep learning methods, RNALens is capable of handling a variety of downstream tasks associated with mRNA expression through fine-tuning. It possesses an exceptionally strong generalization ability and requires less training data for these tasks, making it more suitable for in data-scarce or underexplored domains. In this study, we validate the capability of RNALens in cell- or tissue-specific 5’ UTR function prediction. RNALens consistently outperforms a variety of existing methods, including Optimus, Cao-RF, and other LLM-based models, such as RNAFM MLP, RNABERT MLP, and UTR-LM, in predicting mRNA EL and TE in HEK293T cells, PC3 cells, and muscle tissue, respectively (Figure 1 and Figure 2). This achievement is of significant importance for advancing the field of mRNA therapeutics, where achieving precise expression of therapeutic proteins in specific target cells or tissues is key to improving efficacy and safety [10]. RNALens’s capability to accurately predict cell-specific activity in silico will greatly reduce the experimental burden of screening large 5’ UTR libraries in multiple cell types, thereby accelerating the design process for optimized mRNA drug constructs [13].

### 3.2 Cell-Type Similarities and Regulatory Heterogeneity Revealed by RNALens

RNALens not only excels in prediction accuracy but its predictions also offer some interesting biological insights. The study found a much similar 5’ UTR function on mRNA EL and TE between HEK293T cells and PC3 cells than between either of these two cell lines and muscle tissue (Figure 3 and Table 2). This observation is highly consistent with the biological backgrounds of these cells and tissue. HEK293T cells and PC3 cells both originate from the urogenital system and may share more similar intracellular environments and translation regulatory mechanisms, such as possessing similar active RBP or miRNA expression profiles. In contrast, muscle tissue may differ significantly from the former two in both cellular composition and physiological function. More importantly, muscle tissue itself is heterogeneous, containing multiple cell types [49], so its 5’ UTR library experimental data may reflect an average or combined effect of these different cell types. RNALens’s capability to capture these subtle but biologically significant intercellular differences indicates that the model has learned not just universal sequence features but also regulatory signals related to specific cellular environments. This further implies that RNALens, without explicit information on RBP or miRNA binding sites as input, may have indirectly learned the differences in the activity profiles of these trans-acting factors in different cell types and their ultimate impact on 5’ UTR function.

### 3.3 RNALens as a Virtual Laboratory to Learn the Function of RNA Sequences

The same set of sequences cannot be measured by RNA-seq and Ribo-seq simultaneously since mRNA fragments without ribosome binding will be degraded by RNase before Ribo-seq [50]. However, our models provide virtual experimental environments to predict these two values with high accuracy at the same time. The development of RNALens indeed provides researchers with a potential and powerful in silico platform for rapid, large-scale prediction and characterization of 5’ UTR. This computational approach is orders of magnitude faster and significantly cheaper than performing equivalent wet-lab experiments [13].

We are expecting that the model is able to generate testable scientific hypotheses. For example, providing predicted 5’ UTR variants that are able to maximize EL in a specific cell type while minimizing its expression elsewhere. These computational predictions may then guide subsequent experimental designs, prioritizing the validation of the most promising candidate sequences. Furthermore, by analyzing the sequence features that drive RNALens predictions with the aid of explainable AI techniques, researchers can gain deeper insights into the “regulatory grammar” of 5’ UTRs in different cellular contexts [13], thereby uncovering new regulatory mechanisms.

### 3.4 Limitations of RNALens and Future Directions

Despite the encouraging results achieved by RNALens, there are still some limitations in this study. First, in this study, we used a 100-nt 5’ UTR library [45] for training and validation, while the length of the 5’ UTR is variable [41]. Second, although RNALens makes accurate predictions, understanding the underlying rationale behind its specific predictions remains challenging, which is an inherent limitation of many deep learning models. Third, currently, the model is primarily tested and validated in HEK293T, PC3 cells, and muscle tissue, where its effectiveness for a broader array of primary cells or tissue types is expected to be evaluated in future studies.

## 4 Methods

### 4.1 Overview of RNALens

To enable rich and context-aware representation learning for 5’ UTRs, we develop RNALens, a novel RNA foundation model with a Transformer encoder architecture. The overall architecture of our approach is presented in Figure 4. The model is pre-trained in two stages. The first stage involves standard Masked Language Modeling on general multi-species genomes. At the second stage, the model is trained on 5’ UTR sequence data. As a result, the model learns generalizable regulatory grammar while refining sensitivity to features of 5’ UTR sequences. Subsequently, RNALens is fine-tuned on a range of down-stream tasks related to 5’ UTR regulatory function.

**Figure 4:**
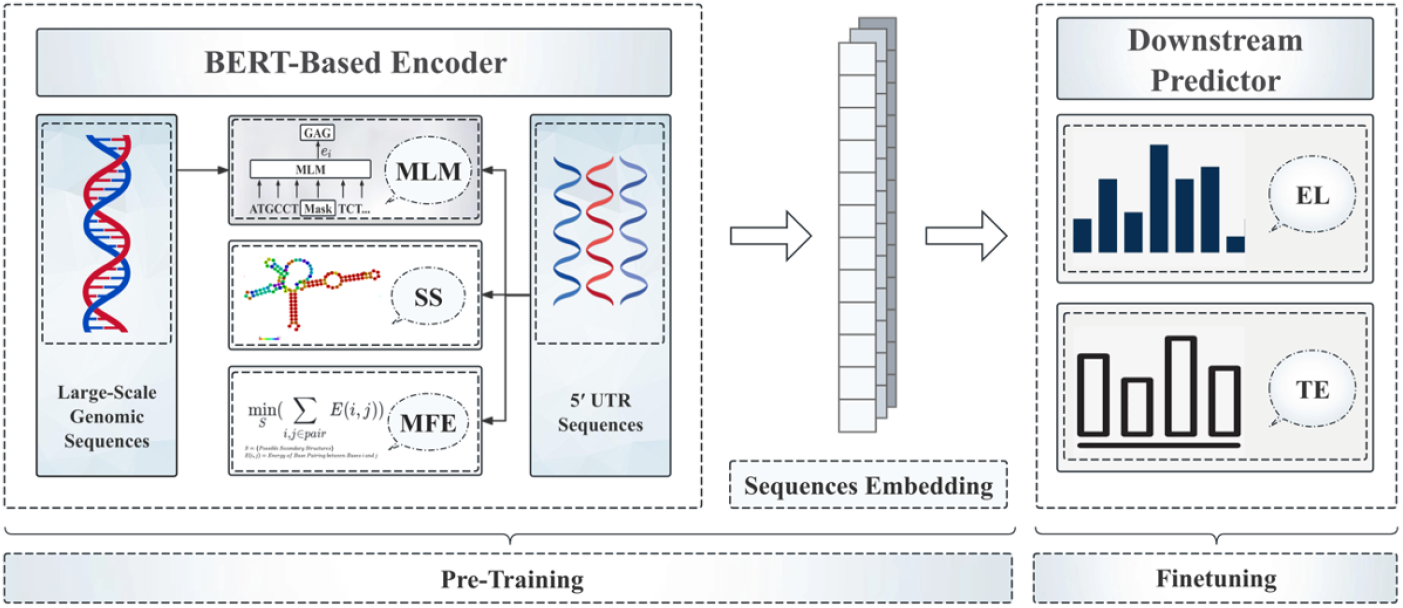
The overall architecture of our approach.

### 4.2 Datasets

The first-stage pre-training uses genome-scale sequences, including the human reference genome and multispecies genomic data, consistent with the datasets previously used to pretrain DNABERT2 [34] and DNAZen [39]. The second-stage pretraining data is a curated collection of 5’ untranslated region (5’ UTR) sequences, designed to capture regulatory patterns specific to UTRs, following UTR-LM [41].

For the 5’ UTR-specific data, we collect unlabeled 5’ UTR sequences from five species—human, rat, mouse, chicken, and zebrafish—sourced from the Ensembl databases [51], as well as endogenous human 5’ UTR sequences previously analyzed by Cao et al. [45]. These human sequences are derived from three biologically distinct sources: HEK293T cells, PC3 cells, and skeletal muscle tissue. Raw sequence data are preprocessed using protocols established by Sample et al. [28] and Cao et al. [45], with strict filtering to retain only high-quality and well-defined 5’ UTRs.

The final pretraining dataset includes 214,349 unlabeled 5’ UTR sequences from Ensembl and an additional 14,410, 12,579, and 1,257 endogenous human UTR sequences from HEK293T, PC3, and muscle tissue, respectively.

### 4.3 Pretraining

The architecture of RNALens consists of a BERT-based encoder block and a downstream predictor block. Each genomic or 5’ UTR sequence of length *L* is input as a series of nucleotides (e.g., ‘A’, ‘G’, ‘C’, ‘T’) with a special [CLS] token prepended. These tokens are then processed using a byte pair encoding (BPE) tokenizer, which merges frequently co-occurring nucleotide patterns into variable-length subword units. The resulting token sequence is embedded and passed through the encoder to produce contextualized representations for each position.

Pretraining is performed in two sequential stages. In the first stage, RNALens is exposed to large-scale genomic sequences—including human and multispecies genomes—to capture global and evolutionarily conserved regulatory features. This stage follows the standard masked language modeling (MLM) objective, in which 15% of input tokens are randomly masked and the model is trained to reconstruct the original nucleotides.

In the second stage, RNALens follows the pretraining strategy of UTR-LM and undergoes task-specific pretraining on curated 5’ UTR sequences. It uses the MLM loss for masked nucleotide reconstruction as well as two auxiliary tasks to integrate structural and thermodynamic information. The first is a secondary structure (SS) prediction task, where dot-bracket notation labels (computed by ViennaRNA) are predicted at masked positions using an MLM-style objective. The second is a minimum free energy(MFE) regression task, where the model predicts the minimum free energy of the input sequence based on the [CLS] token representation. This objective is optimized with mean squared error loss and is motivated by the established correlation between MFE and translation efficiency (TE).

### 4.4 Downstream Finetuning

Protein production is regulated at multiple levels, among which mRNA expression level (EL) and translation efficiency (TE) play distinct yet complementary roles. EL, measured by RNA-seq (e.g., RPKM), reflects the steady-state abundance of mRNA transcripts, and is determined by both transcriptional activity and mRNA degradation. In contrast, TE, typically defined as the ratio of Ribo-seq to RNA-seq signal, captures translation dynamics and ribosome recruitment. Predicting EL and TE from 5’ UTR sequences offers insight into UTR-mediated regulation. Both are critical determinants of the final protein output of an mRNA, making them essential for guiding the rational design of mRNA-based therapeutics.

To fine-tune RNALens for these tasks, we utilize 5’ UTR datasets [45], derived from three distinct human cell or tissue types: HEK293T, PC3, and skeletal muscle. These datasets contain a total of 41,446 unique 5’ UTR sequences, each associated with paired measurements of EL and TE. Following the experimental design of Cao et al., we fixed the UTR length to 100 nucleotides.

For each cell or tissue type, we fine-tune separate RNALens models for EL and TE prediction using the pretrained weights as initialization. The human endogenous 5’ UTR sequences are processed with the same BPE tokenizer as in pretraining, and split into training, validation, and test sets. Each model is trained for 300 epochs, and performance is evaluated on the test set using the Spearman correlation coefficient between predicted and experimentally measured values.

To assess the cell-type specificity and generalization capacity of RNALens, we further evaluate cross-cell performance by applying a model trained on one cell type (e.g., HEK293T) to the test data from another cell type (e.g., PC3 or muscle). This allows us to examine whether the model had captured cell-specific regulatory features or learned transferable representations across cellular contexts.

## Footnotes

1 The code and models are available at https://github.com/oomics/RNALens.

2 The library comprises approximately 12,000 100-nt 5’ UTR sequences with corresponding RNA-seq data and Ribo-seq data in PC3 cells (human prostate cancer cell line), HEK293 cells (human embryonic kidney cell line) and muscle tissues [41].

